# Resource competition controls the length-dependence of synaptic protein synthesis

**DOI:** 10.64898/2026.10.30.763010

**Authors:** Patricia Rubisch, Sofia B. Raak, Judith A.H. Wodke, Melanie I. Stefan, Cian O’Donnell

**Author notes:** Senior author (Melanie I. Stefan, Cian O’Donnell).

## Abstract

Synaptic plasticity is dependent on the availability of specific proteins at the synapse with distinct requirements for potentiation and depression. Specifically, experimental evidence suggests an upregulation of short proteins and a downregulation for long proteins during long-term depression (LTD). The mechanism of this length-dependent protein synthesis regulation is yet unknown. In this work, we propose an computational model of ribosomal regulation of protein synthesis than can reproduce the length-dependent shift in protein expression seen during ribosomal elevations. By analysing the model dynamics, we discover that this effect is due to a shift from initiation-restricted to elongation-restricted protein synthesis. This works offers a novel explanation for the loss of ribosomal control of protein synthesis due to saturation in pathological ribosome enriched conditions. This new theoretical model of ribosomal regulation of protein synthesis is capable of explaining length-dependent shifts in protein synthesis.

**Author Summary:** At synapses, the connection between neurons in the brain, the amounts of specific proteins determine the strength of communication between the neurons. The process of translation followed by protein synthesis provides the cell with the proteins to uphold the synaptic strength. A change in the translational profile and therefore a change in amount of the synthesised proteins can result in synaptic plasticity, the strengthening or weakening of the synapse. The required changes in the synaptic translation profile are non-trivial. With an increase in ribosomes, needed during translation initiation, long proteins are downregulated while short proteins are upregulated. The mechanism of this length-dependent translation profile shift is yet unexplained. In this computational study, we investigate resource abundances during translation as a possible explanation. We introduce a novel, minimal model of translation consisting of three phases: initiation, elongation and degradation. We assume that initiation and elongation are subject to resource restrictions. The interplay between resource availability in the model explains the length-dependent shift in translation profiles. An increase in ribosomes allows the system to dynamically switch from initiation-restricted to elongation-restricted dynamics. Furthermore, the model offers a novel explanation for the loss of ribosomal control in pathological conditions. In highly ribosome enriched conditions, the dynamics reach saturation resulting in translation profiles which are insensitive to changes in ribosomal availability. In summary, our model shows that resource competition during initiation and elongation is necessary for the length-dependent regulation of the translation profile.

## Introduction

Synaptic proteins are central for maintaining synaptic function. Due to turnover and protein degradation, the synapse needs a steady supply of synthesised proteins^1,2^. The change of synaptic strength, Long-Term Plasticity, requires specific changes in the local protein abundances^3–12^. To express Long-Term Depression (LTD) by weakening the synapse, AMPA receptors need to undergo endocytosis requiring metabolic proteins. In contrast, the expression of Long-Term Potentiation (LTP), strengthening the synapse, requires a higher amount of newly synthesised AMPA receptors. Thus, dysregulation of translation and protein synthesis at synapses can hinder synaptic learning leading to memory loss or learning disability on the behavioural level^13,14^.

Protein translation can be controlled by ribosome abundance^15,16^. The more ribosomes are present, the higher the likelihood one will bind to an mRNA. Thus, changes in ribosome abundance act globally and and exert proteome-wide effects. Surprisingly however, the observed effects are non-trivial: an increase in ribosome numbers does not increase the protein yield uniformly^17,18^. Instead experimental studies report a length-dependent effect. An increase in ribosome abundance will lead to an increase in translation of short mRNAs and a decrease in translation of long mRNAs^19,20^. The underlying mechanism of this length-dependent regulation is currently unknown.

Proteome-wide effects due to an aberrant translation regulation are also present in *Fmr*1^−*/y*^ mice, a genetic model of Fragile X Syndrome^21,22^. In *Fmr*1^−*/y*^, higher ribosomal availability appears due to an increase in ribosomal abundance or reduction of ribosomal stalling^19,23,24^. Interestingly, Seo et al. ^19^ report similarities between the length-dependent up-/downregulation of synaptic proteins between the *Fmr*1^−*/y*^ mouse model and wild-type mGluR-induced LTD. Both *Fmr*1^−*/y*^ and mGluR-LTD proteomes show an upregulation of short proteins associated with with metabolic function while long synaptic proteins are downregulated. Additionally, ribosomal protein abundance is also increased in both conditions. In mGluR-LTD this increase is caused by the stimulation of ribosynthesis^14,25^, increasing the ribosomal abundance dynamically.

In this work, we explore the effect of resource availability and competition in minimal mathematical models of protein translation using systems of Ordinary Differential Equations (ODEs). Our goal is to discover regimes of competition in the model that are sufficient to reproduce the experimentally observed length-dependent gene expression changes following changes in ribosome abundance. We systematically consider the effect of competition for two limited resources: 1) ribosome availability during initiation, and 2) amino acid availability during elongation. We identify that the requirements for reproducing the experimental data are only met when initiation and elongation are dependent on a shared, limited resource. Therefore, our model predicts that ribosomes and amino acids are competitive resources. Changes in their abundances can regulate protein synthesis length-dependently as necessary for the expression of synaptic plasticity. In the future, experimental data systematically manipulating the resource abundance *in vitro* is needed to enable model-fitting and quantitative predictions.

## Results

We aim to assess various hypothetical explanations for the length-dependent ribosomal regulation of protein expression (Figure 1). We hypothesised that the phenomenon could be explained by either or both of ribosome competition at translation initiation, or amino acid competition during elongation. We assess this using different versions of a simple mathematical model to rule in or out the various resource competition combinations.

**Figure 1.**
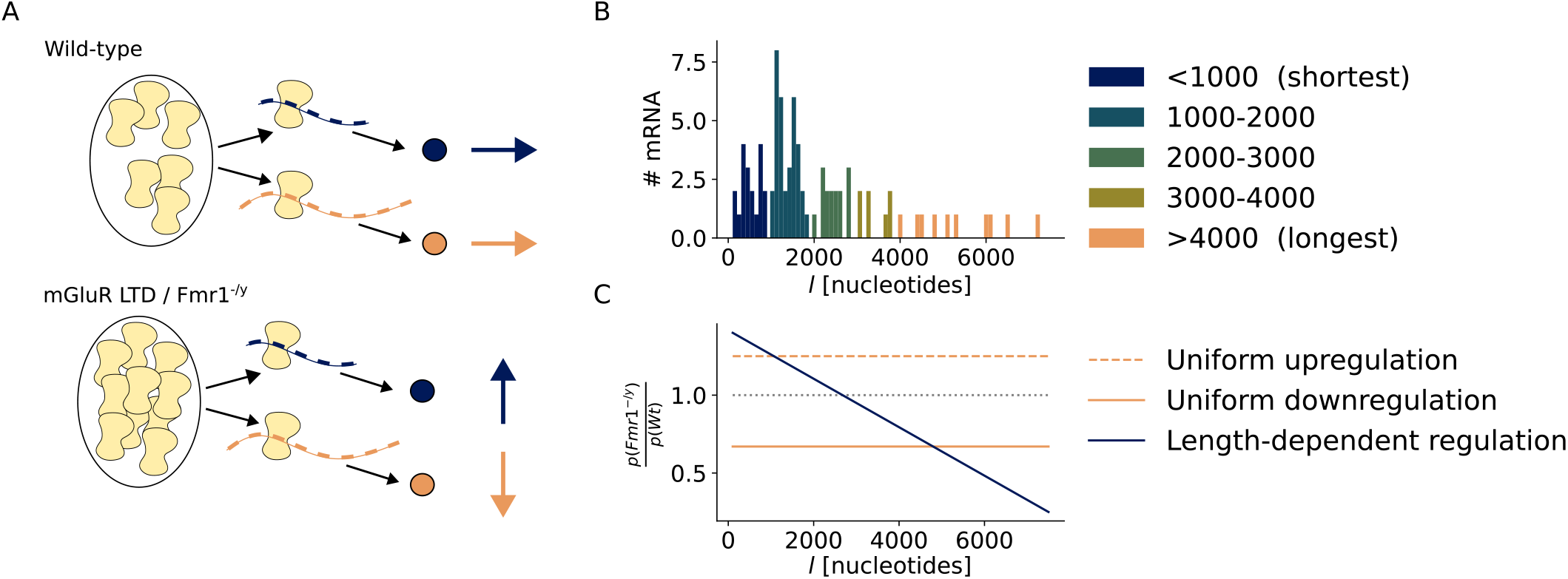
Translation profile shifts and changes in ribosomal abundance in *Fmr*1^−*/y*^ model correlates with mGluR LTD induced changes in wild-type. **A**: Illustrative summary of wild-type and high ribosome abundance states. **B**: Considered mRNA population in number of amino acids characterised by mRNA length *l*. **C**: Visualisation of the ribosomal regulation hypotheses. The naive assumption of uniform, length-independent regulation is shown in yellow. The experimentally observed length-dependent regulation of the translation profile is shown in blue. Depicted is the ratio of the protein yield between the high ribosome abundance state (basal *Fmr*1^−*/y*^) and the low ribosome abundance state (basal wild-type).

### Minimal ODE description of translation allows for large population analysis

Protein translation is described in three steps: initiation, elongation and termination. We allocated a dedicated rate parameter *α* for the initiation step. But to keep the model simple, we combined the elongation and termination steps into one process governed by a single rate parameter *β*. We study a model of protein synthesis and degradation where during initiation, free mRNAs *x*_*i*_ bind to ribosomes *r* and form an active complex 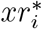, where *i* indexes the particular mRNA type. During elongation and termination, the active complex sequesters charged tRNAs loaded with the amino acid *y* and synthesizes protein yield *p*_*i*_. This protein can then spontaneously degrade at rate *γ* to replenish the amino acid pool. Note this simplified model aggregates elongation and termination into one step, and abstracts the degradation and resource recycling process. In the following the aggregated elongation and termination step is referred to simply as elongation for brevity.

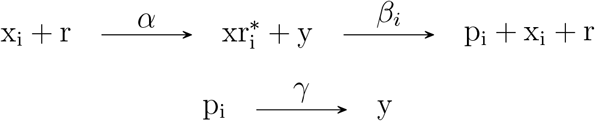

This system has two finite resources: ribosomes *r* and amino acids *y*. If ribosomes are scarce, translation initiation rates will be restricted. In contrast if amino acids are scarce, translation elongation rates will be restricted. We considered three different resource restriction scenarios:

1. the resource is unlimited and homogenously distributed
2. the resource is limited and spatially restricted resulting in multiple individual resource pools for each mRNA type
3. the resource is limited and spatially unrestricted and therefore shared between mRNA types

Representing all possible combinations of two resources under three availability conditions results in 3 *×* 3 = 9 minimal models. A summary of the resource limitations along the mathematical definition of the resources within the specific model is provided in Table 1.

**Table 1.**
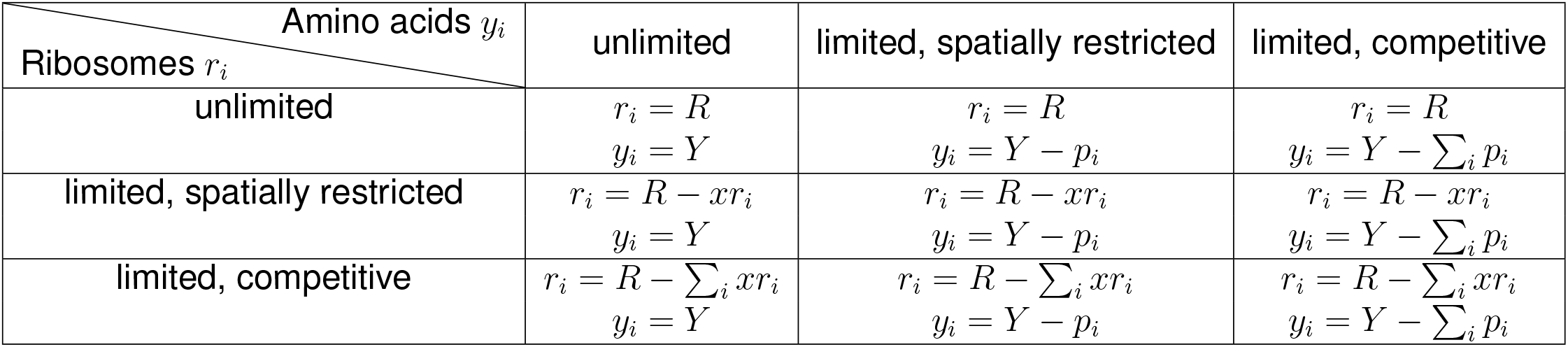
Mathematical description of the resource assumption for all possible nine combinations for ribosome and amino acid availability.

| Amino acids *y*<sub>*i*</sub><br>Ribosomes *r*<sub>*i*</sub> | unlimited | limited, spatially restricted | limited, competitive |
|---|---|---|---|
| unlimited | *r*<sub>*i*</sub> = *R*<br>*y*<sub>*i*</sub> = *Y* | *r*<sub>*i*</sub> = *R*<br>*y*<sub>*i*</sub> = *Y* − *p*<sub>*i*</sub> | *r*<sub>*i*</sub> = *R*<br>*y*<sub>*i*</sub> = *Y* − ∑<sub>*i*</sub> *p*<sub>*i*</sub> |
| limited, spatially restricted | *r*<sub>*i*</sub> = *R* − *xr*<sub>*i*</sub><br>*y*<sub>*i*</sub> = *Y* | *r*<sub>*i*</sub> = *R* − *xr*<sub>*i*</sub><br>*y*<sub>*i*</sub> = *Y* − *p*<sub>*i*</sub> | *r*<sub>*i*</sub> = *R* − *xr*<sub>*i*</sub><br>*y*<sub>*i*</sub> = *Y* − ∑<sub>*i*</sub> *p*<sub>*i*</sub> |
| limited, competitive | *r*<sub>*i*</sub> = *R* − ∑<sub>*i*</sub> *xr*<sub>*i*</sub><br>*y*<sub>*i*</sub> = *Y* | *r*<sub>*i*</sub> = *R* − ∑<sub>*i*</sub> *xr*<sub>*i*</sub><br>*y*<sub>*i*</sub> = *Y* − *p*<sub>*i*</sub> | *r*<sub>*i*</sub> = *R* − ∑<sub>*i*</sub> *xr*<sub>*i*</sub><br>*y*<sub>*i*</sub> = *Y* − ∑<sub>*i*</sub> *p*<sub>*i*</sub> |

In the model, an individual mRNA species *x*_*i*_ is defined by its elongation rate *β*_*i*_. The longer the transcript *x*_*i*_, the more time a single ribosome needs to traverse the full transcript length. To capture this effect we assume elongation rate *β*_*i*_ is inversely proportional to the length *l*_*i*_ of the mRNA species. Due to the focus on the length effect, which is absorbed by the definition of the elongation rate *β*_*i*_, we assume that initiation rates are homogenous. Furthermore, the model is agnostic about the contribution of secondary structures or codon-specific elongation rates.

### Ribosome regulation of protein synthesis depends on amino acid abundance

In the following, we systematically test all model variations for their capacity to reproduce the length-dependent shift of the translation profile in ribosome enriched conditions. We use a compact description defined by the difference in protein yield ratios as model evaluation metric. We identify the elongation resource competition for amino acids as a necessary condition for reproducing the experimentally observed pattern.

To evaluate whether the different model variations are able to reproduce the length-dependent up-/downregulation, we use the ribosome-dependent change in protein yield ratios *ρ*:

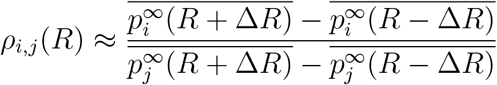

*ρ* is calculated by approximating the protein yield gradients for a subpopulation of short proteins *p*_*i*_ and a subpopulation of long proteins *p*_*j*_ by computing the population averaged slopes of the respective populations. Thus, *ρ*_*i,j*_(*R*) *<* 0 only if exactly one of the approximate population gradients is negative (more details provided in Methods). This means, that the ratio can only be negative if either the long proteins increase with increasing ribosome numbers while the short proteins decrease or the long proteins decrease with increasing ribosome number while the short proteins increase. Therefore, a negative ratio indicates a length-dependent shift characterised by the suppression of translation of one of the groups.

To assess the suitability of all nine possible model variations, we compute *ρ* across a wide range of initiations rates *α* and elongation rates *β*. The results shown in Figure 2 indicate that a competition for the elongation resource is a necessary condition for the desired pattern to be reproduced. This is shown by the change from positive values of *ρ* (red) to *ρ <* 0 (blue) in the heat maps. Any other model variation is characterised by a plot which only produces *ρ >* 0 across the parameter ranges. These models predict a regulation in the same direction for long and short mRNAs.

**Figure 2.**
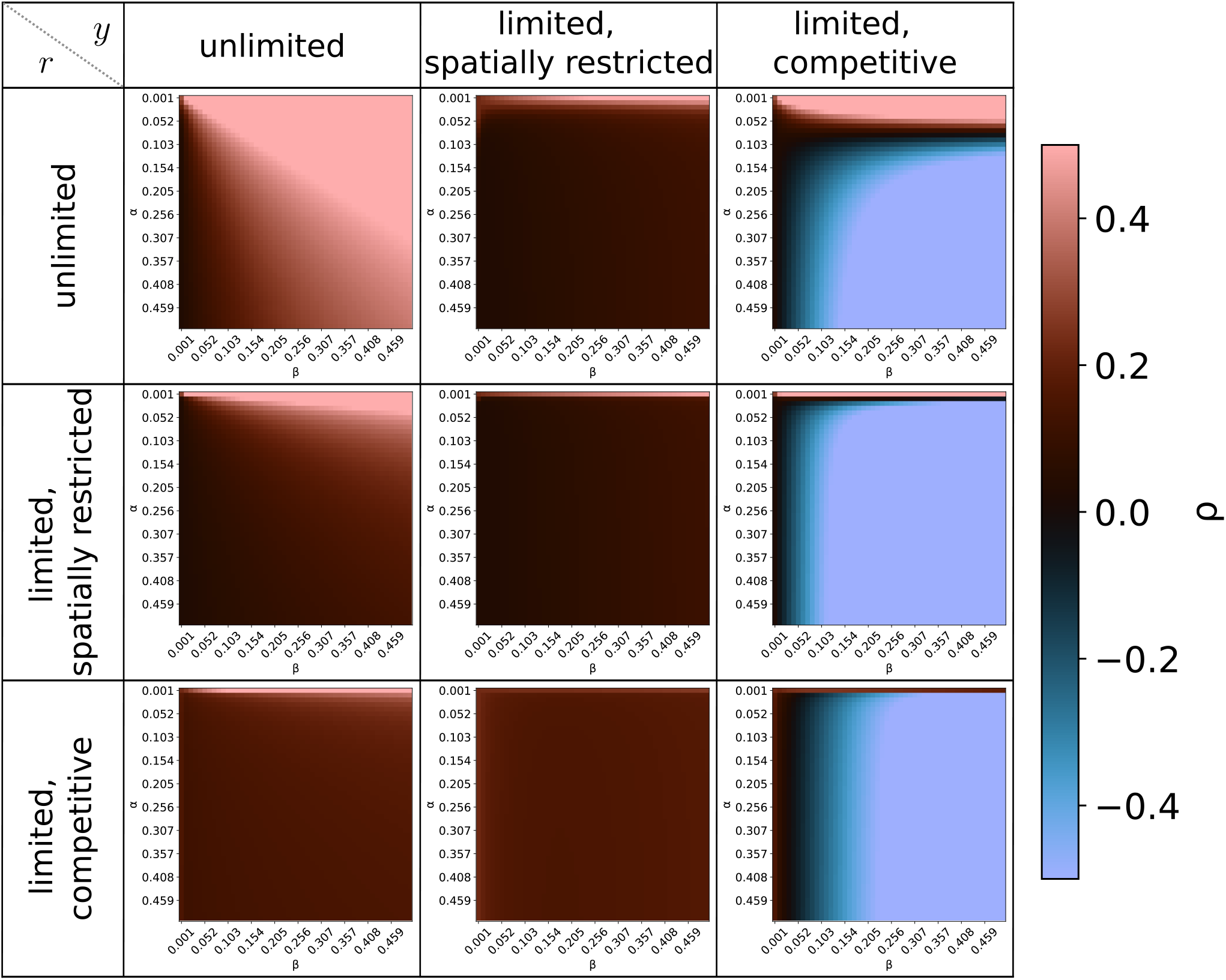
Influence of parameter variation on competition measurement *ρ* identifies competitive amino acid resource as necessary condition to reproduce experimental data. The ratio of the protein yield change *ρ*_*i,j*_ is shown for all nine model variations. Model variations are indicated by row and column. The heatmaps show the dependence of *ρ*_*i,j*_ on the initiation rate *α* (y-axis) and the elongation rate *β* (x-axis). The value of *ρ* is computed at *R* = 75 with Δ*R* = 25 with a population split *p*_*i*_ = *i* : *l*_*i*_ *>* 2*k* (long) and *p*_*j*_ = *j* : *l*_*j*_ *<* 2*k* (short). The results are shown as a diverging colourmap with positive measures of *ρ >* 0 shown as red hue and *ρ <* 0 depicted as shades of blue.

The three candidate models do not show the same relationship between *α, β* and *ρ*. They show similarities as the negative correlation between *ρ* and *β*. The higher the elongation rate, the stronger is the measured competition shown by a more negative value of *ρ*. The candidate model variation which assumes that ribosomes are an mRNA species-specific resource (“limited, spatially restricted”) shows a wider range of initiation rates *α* for which the model predicts a uniform ribosomal regulation.

To further explore the differences driven by the initiation resource condition, we explore the ribosomal dependence of the protein yield using our standard parametrisation (see Methods for details). Comparing the protein yield and *ρ*-curves (Figure 3A and B) shows that the ribosomal resource assumption determines the range over which ribosomal regulation occurs. All models show a *ρ*_*i,j*_ *<* 0 after a critical ribosome abundance *R*^*^ and a saturation of *ρ*_*i,j*_ indicating that while the protein yields change, the relative amount of long to short proteins do not change (Figure 3B). However, they differ in the range over which the *ρ*_*i,j*_ saturation point is approached. When ribosomal abundance simply scales the initiation rate, the effect range is small. Saturation is reached for most gradient ratios when *R*^*^ *<* 1. In contrast, when ribosomes are modelled as mRNA-specific finite pools, the range is significantly broader. In comparison, this model variation shows the broadest range for approaching the *ρ*_*i,j*_ saturation point. Assuming that ribosomes are a finite resource which is shared between translating transcripts the protein yields reach their saturation point at a comparably intermediate ribosome abundance. The model variations also show distinct differences in the protein yield behaviour which can be characterised using 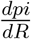 (Figure 3C). The gradients also show the differences in effect ranges as described for *ρ*_*i,j*_.

**Figure 3.**
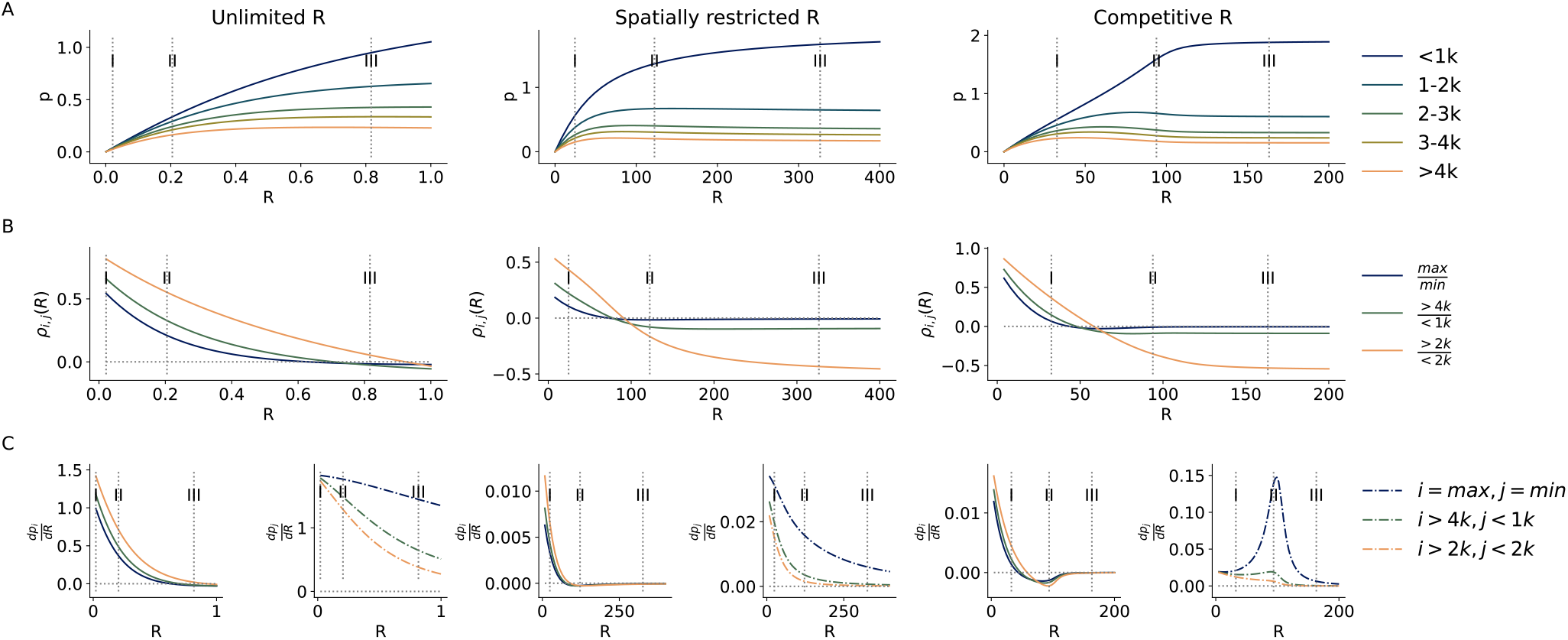
Shared ribosomal resource is necessary for active changes in length-dependent protein yield at intermediate ribosome abundance. Results are shown per ribosomal resource assumption (column). **A**: Protein yield. **B**: *ρ*_*i,j*_ for three different subpopulation compositions. **C**:Approximated gradients of subpopulations shown in B.

In all models, long transcripts, *l >* 1*k*, show three distinct regulatory regimes: Increasing ribosome abundance leads to an I. increase in protein yield. The gradients are positive. This if followed by a II. reduction in protein yield, resulting in negative gradients. If ribosomal abundance is sufficiently large, a III. saturation regime is reached in which the gradient is near 0. Short transcripts, *l <* 1*k* can be characterised by their positive gradients. They do not show a ribosome abundant sign switch: An increase in ribosome abundance causes a growth in the protein yield. However, model variation which do not assume the ribosome abundance to be shared between mRNA species shows monotonously increasing gradients. When ribosomes are assumed to be a competitive resource, the gradient reaches a peak at intermediate ribosome abundance. This indicates that the dynamics contributing to the length-dependent protein yield change when both ribosomes and amino acids are a restricted and competitive resource.

### Ribosomal regulation and saturation regime is defined by effective initiation and elongation rates

The protein yield is dependent on the effective initiation and elongation rates which are scaled by the ribosomal abundance and amino acid abundance respectively. Therefore, the maximally available resource abundance determines the dynamics and the saturation regime of the protein synthesis. Analysing the relationship between the effective rates and their drain on their respective resources, identifies the model which assumes both ribosomes and amino acids as restricted and shared resources as the best approximation of the experimental observations.

In the following, we define the effective initiation and elongation rates as the net increase of the actively translating ribosome-mRNA complexes and the net increase in protein yield, respectively:

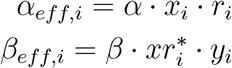

The comparison of these two quantities allows us to identify the driving quantity behind the protein-yield dynamics. If *α*_*eff,i*_ *> β*_*eff,i*_, elongation restricts the efficient translation of the initiated complexes, the species is elongation-restricted. Alternatively, if *α*_*eff,i*_ *< β*_*eff,i*_, initiation is limiting the amount of actively translating complexes, the species is initiation-restricted. Furthermore, the steady-state in this system at which protein yield is stable is defined by *α*_*eff,i*_ = *β*_*eff,i*_ (see Methods). Therefore, we can characterise the different dynamical regimes using *α*_*eff,i*_ *− β*_*eff,i*_ as a measurement of the restricting condition.

As Figure 4 shows, the dynamics in the competitive ribosome assumption model differ from the other variations in which ribosomes are not shared. The dynamics of short mRNA transcripts undergo a change from being initiation-restricted to being elongation-restricted at high ribosome levels. That this change only occurs in this model is due to the competition for the ribosome resource.

**Figure 4.**
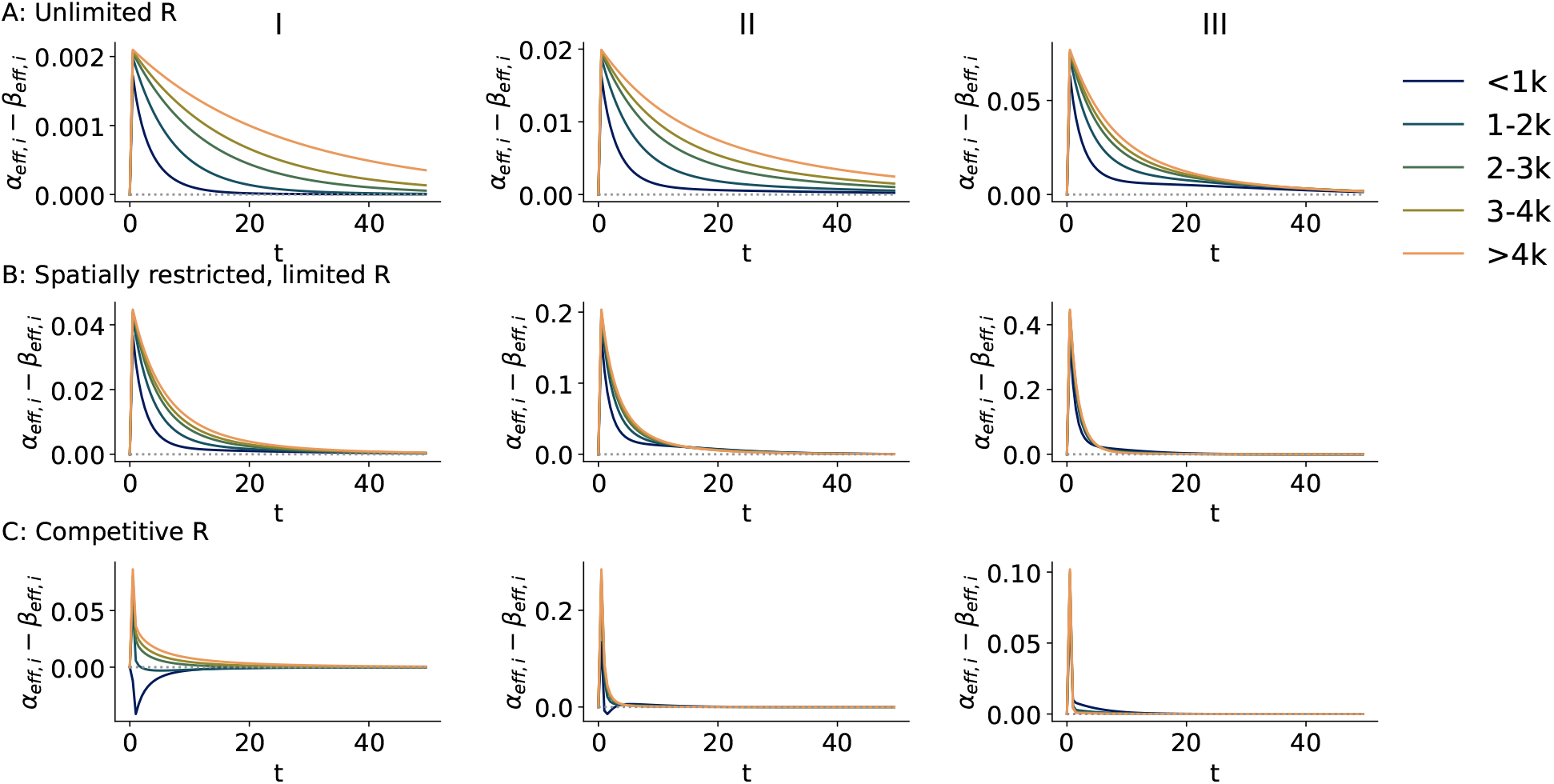
Shared ribosomal resource results in a change from initiation to elongation restriction for short mRNAs at intermediate ribosomal abundance. I, II and III corresponds to the indicated ribosomal concentrations in Figure 3. The time course shows the difference of effective rates *α*_*eff,i*_ *− β*_*eff,i*_ for the five length-dependent mRNA population splits. **A**: Unlimited ribosome model. **B**: Limited, spatially restricted model. **C**: Shared ribosome model.

Figure 5 depicts the freely available resources in each condition over the first 50 time units. When ribosome abundance scales the effective initiation rate (”Unlimited R”), the level is not affected by the protein synthesis as by definition. The ribosomal abundance determines the max of *α*_*eff,i*_ but all effective initiation rates are equally scaled. Therefore, the effect of *R* is static over time and linear over the ribosomal abundance range. When ribosomes are spatially restricted but mRNA transcript-specific, their scaling effect is minimally delayed. However, as only *x*_*i*_ drains *r*_*i*_, the free ribosome abundance is completely used up within a short time window. Effective *r* also scales *α*_*eff,i*_. All species of the mRNA population are equally effected by this scalar. Therefore, *R* only determines the maximal effective initiation rate, but does not effect the interplay between effective initiation and elongation at different rates. Disregarding the overall ribosome abundance, the system is always in an elongation-restricted regime. Therefore, the total protein yield of any mRNA transcript is predominantly determined by the amino acid abundance while the ribosomal abundance reduces to a scaling factor. Due to the length-dependence of the elongation rate *β*_*i*_, we observe a length-dependent protein yield. However, the shift of the translation profiles is driven by the more effective translation of the individual species, not the ribosomal abundances by itself.

**Figure 5.**
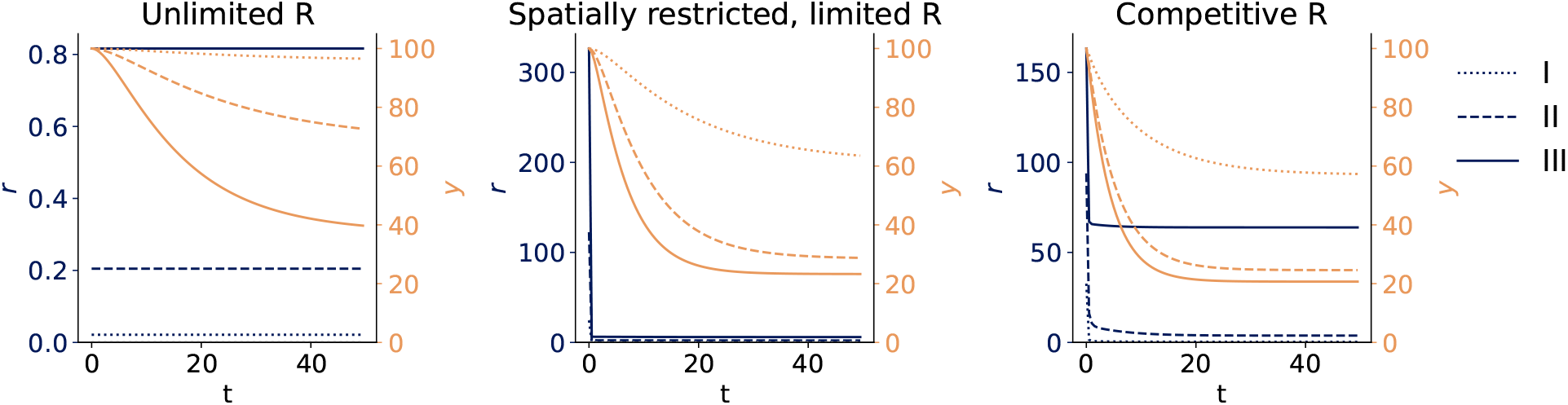
Ribosomal resource depletes disregarding its maximal concentration when ribosomes are modelled as a non-competitive resource. Time course of resource depletion of ribosomes *r* (blue) and amino acids *y* (yellow).

It is clearly visible in Figure 5, that while the drain on the ribosome abundance stabilises and finds its steady-state quickly, the drain differs between conditions. *R* does not simply scale the effective initiation rate linearly between conditions. At low ribosome conditions, the pool is completely drained which agrees with observations that short proteins are initiation-restricted (Figure 4CI). When ribosome abundance increases, the system shifts towards elongation restriction for all mRNA transcripts. At saturation (III), the ribosome abundance is big enough, that *α*_*eff,i*_ can reach its maximum disregarding the species and associated elongation rate. At intermediate ribosome abundances, the system undergoes a positive feedback loop between effective initiation rate and effective elongation rate for short proteins (II), visible in a slower dynamic towards the steady-state of the free ribosomes (see 5).

In these instances, the initial *α*_*eff,i*_ is set by the ribosome abundance for all mRNA species. For short proteins, for which *β*_*i*_ is comparatively large, *β*_*eff,i*_ is also large. This means that all *r*_*i*_ used for the translation of short mRNA are returned to the pool of free ribosomes quicker. This increases *α*_*eff,i*_ for all mRNA due to the length-independence of *α*. Due to the increase in 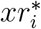, *β*_*eff,i*_ increases and short transcripts are more quickly released from the ribosomes again. If the increase in free ribosomes is significantly big, this can cause the *α*_*eff,i*_ *> β*_*eff,i*_ also for short mRNAs. This interplay continues until the free ribosome abundance stabilises.

In summary, when ribosomes are a shared commodity, short proteins can more efficiently translate at intermediate ribosome abundances. Due to a positive feedback-loop between effective initiation and elongation rates, the synthesis of short proteins shifts from an initiationrestricted to an elongation-restricted translation regime.

### Basal *Fmr*1^*−/y*^ protein synthesis lies in the saturation region

We have identified the necessary resource assumptions which need to be fulfilled in order to observe active ribosomal regulation of protein synthesis. Both initiation and elongation need to be subject to resource restriction. In the following, we elaborate on the biological implications of the model predictions. We identify possible regions for the pathological basal state of *Fmr*1^−*/y*^, as well as the directional shifts and further predictions of translation shifts during plasticity induction proposing a unifying theory of ribosomal regulation of neural protein synthesis (Figure 6).

**Figure 6.**
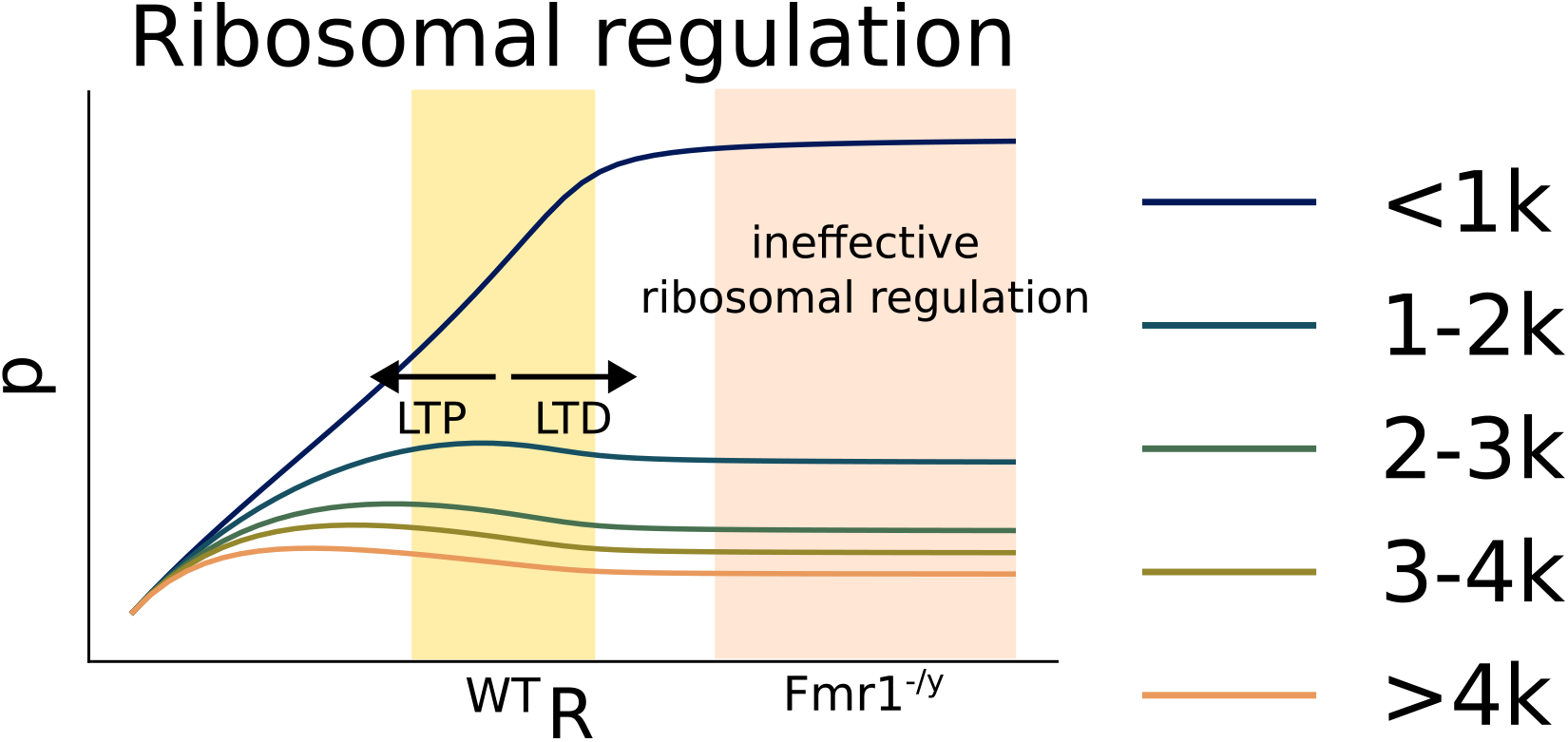
Model indicating saturation (*Fmr*1^−*/y*^) and effective ribosomal regulation (WT) regimes. Reduction of ribosomes in the effective regulation regimes leads to a decrease in short proteins and an increase in long proteins. Ribosome enrichment leads to the opposite pattern: Long proteins decrease and short proteins increase. Ribosomal depletion or low abundance leads to a uniform, length-independent regulation.

The candidate model assuming both ribosomes and amino acids as shared resources for the translation of mRNA transcripts predicts a critical point *R*^*^ at which saturation of the system is achieved (Figure 6 orange background). If *R > R*^*^, the translation profile at *R* will not differ from the profile at *R*^*^. Seo et al. ^19^ show that mGluR-dependent ribosynthesis in the *Fmr*1^−*/y*^ model, an increase in ribosomal proteins in the neuron, does not causes significant changes in the translational profile. Therefore, our model identifies the saturation regime as the pathological state of *Fmr*1^−*/y*^. Due to the sensitivity of the translational profile with regard to ribosynthesis after LTD induction, the wildtype basal and mGluR-LTD states are predicted to be situated at intermediate ribosomal abundances (yellow background). In this range, increasing the ribosomal concentration has significant effect on the protein translation profile and the ratio between long and short proteins can be regulated. Furthermore, our model proposes that the proteome changes needed for increasing the synaptic strength after LTP induction can also be caused by manipulating ribosomal abundance. A reduction of ribosomal abundance upregulates long proteins, while short proteins are downregulated. As a consequence the balance between metabolic and synaptic proteins is shifted towards favouring facilitation of synaptic strength and the increase in transmembrane proteins.

## Discussion

This work introduced a minimal ODE model for the description of translation in neurons. We consider both ribosomes and amino acids as shared and finite resources during the process. Thus, initiation and elongation are dependent on their respective resource. We found that not resource scarcity but intermediate levels of ribosomal abundance lead to a competitive effect due to a positive feedback loop between effective initiation and elongation rates of short proteins. It can capture the ribosomal regulation of the translation profile after mGluR-LTD induction and predicts the basal state of *Fmr*1^−*/y*^ neural protein synthesis profiles to lie in the saturation regime where ribosomal regulation is ineffective. Moreover, assuming that LTP is also subject to the ribosomal regulation shifting the translation profile, the model presents a comprehensive theory of ribosomal regulation for the translational effects during plasticity.

### Related work

Previous models of mRNA translation have a higher mRNA specificity. Using ribosomal flow models or TASEP models, one can consider the contribution of codon-specific elongation rates^26**?** – 29^

However, those studies use ribosomal density as a proxy for the protein yield. Furthermore, most models only consider the translation of one mRNA at a time and therefore do not take into account the effect of resource sharing. Raveh et al. ^27^ extended the ribosomal flow model to an interconnected system of multiple mRNA species which drain a common ribosome pool. They were able to demonstrate that the interaction under resource constrains are non-trivial and codon/mRNA-specific. However, they focused their investigation on codon-specific influences while ignoring that elongation is also subject to resource availability in the form of amino acids. Therefore, the model presented here is the first to consider resource susceptibility of both the initiation and elongation phase while at the same time modelling the protein yield explicitly. Moreover, it defines a novel minimal model for the ribosomal translation process with the resulting compact description facilitating understanding of underlying regulatory mechanisms.

### Limitations

While this model presents a compact and minimal description of protein translation, it abstracts away a lot of influences which can also effect resource use during protein synthesis. We assume that the relationship between protein length and elongation rate is strictly monotonous. However, it has been shown that elongation rates are codon-specific. This means that the elongation rate is a product of the number of specific codons and their codon-specific elongation rates^28^. When the mRNA transcripts are known in their composition, a mean elongation rate could be calculated and used for the model which would increase the explanatory power.

On a related note, we currently only know two (unspecific) points within the ribosomal abundance range: the wild-type and *Fmr*1^−*/y*^-mutation basal states. While *in vivo* data is highly desirable, tightly controlling the ribosomal abundance in *in vivo* cells is not currently feasible. Further experimental studies to systematically record translation profiles in dependence to ribosomal and amino acid abundance are needed to fully verify the model. Especially the region of non-pathological basal protein translation is of high interest, as the model predicts this state to be in a highly sensitive regime. The question arises how stable protein synthesis profiles can be guaranteed when the system is sensitive to the exact concentration of ribosomes available. A more extensive body of experimental data points would also make it possible to estimate the mean elongation rates of mRNA transcript translation by fitting the model behaviour to it.

Furthermore, we analyse the system exclusively under static environmental conditions where parameters are fixed in time. These assumptions are not representative of the conditions within the neuron, especially not when considering that ribosomal abundance might be a dynamic regulator of protein translation of plasticity as suggested here. Therefore, future research should explore the dynamics and profile shift predictions under non-static conditions. These could not only arise from active ribosome regulation but also from transcriptional regulation influencing the composition of the mRNA population.

Translation as modelled in this study is perfect. Translation fault, ribosome jamming and detaching before termination is not considered. Similarly circularisation which promotes reinitiation^29,30^ is not modelled. Both of these processes have an effect on effective initiation and elongation rates. They might increase the competitive region due to the favouring of short transcripts for circularisation and the higher probability for faults or ribosomal jams during the elongation of long transcripts. Based on these considerations, the model should predict an exacerbated competition if ribosome jamming or circularisation is included.

We performed a reproducibility audit as suggested by Fitzpatrick and Stefan ^31^ . However, there are currently no standards on how to perform such an audit. Therefore, the exact procedure to follow and possible improvements for future projects were part of the in-person discussion. We hope that the community will join in developing standards and procedures for this kind of important aspect of quality control.

### Summary & Outlook

In this work, we have introduced a novel minimal model for neural protein synthesis. Protein synthesis is especially of interest in neurons due to the demand for a shift in functional protein composition during plasticity induction, which our model reproduces successfully. By systematic exploration of resource conditions concerning ribosomes and amino acids, we demonstrate that shared amino acids as elongation resource are necessary to achieve a length-dependent up-

/downregulation of short/long proteins. Ribosomal competition is needed in order to link the protein synthesis of multiple transcripts and model the active competition for resources. Our findings support studies using interconnected Ribosomal Flow Models^27^ suggesting the active competition for ribosomes during initiation to be a modulator of ribosomal occupancy. We could extend on these insights by studying the influence of ribosomal and amino acid abundance on the composition of translational profiles. This study suggest that ribosomal abundance is a regulator of protein synthesis during LTD in neurons. Focusing on ribosomal abundance in proteome studies after plasticity induction and in further synaptopathologies might provide more insights into the importance of the resource abundance regulation of translation as a functional mechanism in synapses.

## Methods

### Differential systems

We model the chemical reaction system

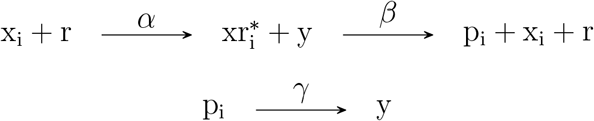

using an ODE system which describes the rate of change for the free mRNA *x*_*i*_, the active complex 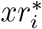 and the translated protein *p*_*i*_. *α* is the initiation rate constant, *β* is the elongation rate constant, and *γ* is the protein degradation rate. As described previously, the amount of free mRNA transcripts is decreased by the initiation process and replenished by the coupled elongation and termination step:

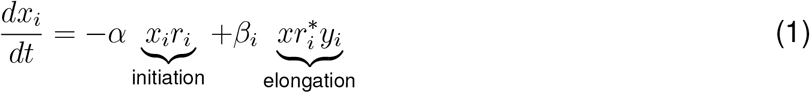

The processes have the opposite effect on the amount of active complexes. Initiation increases the number of active complexes while the elongation/termination reduces the amount:

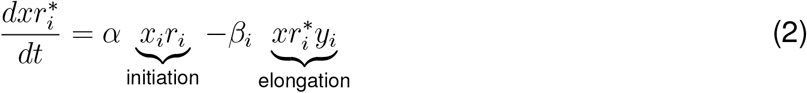

The rate of change of protein is given by the difference between effective elongation and protein degradation:

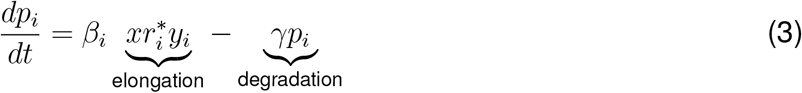

Due to the level of abstraction the ODE system does not account directly for the length of the mRNA. To introduce mRNA-specific length effects, the elongation rate is defined anti-proportional to the mRNA length *l*_*i*_

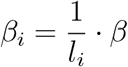

The short the mRNA, the quicker the elongation because less mRNA codon translation steps need to be carried out. If not stated otherwise, the ODE system uses the standard parametrisation noted in Table 2.

**Table 2.**
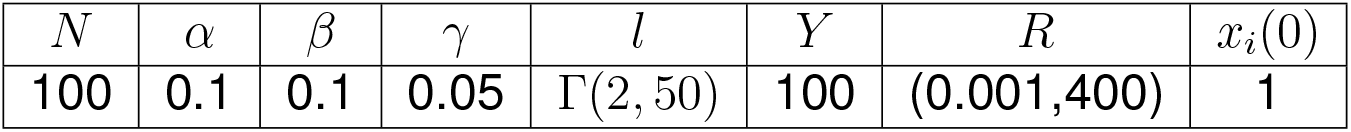
Parametrisation of the model.

The model was simulated for *t* = 200 time steps using the Euler-integration method with an integration time-step of *dt* = 0.001.

### Steady-state and effective rates of the ODE system

If not otherwise indicated in the plot, all results are shown at the approximate steady-state. The steady-state of the system is defined by:

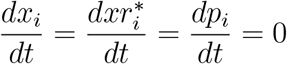

Thus, the following equations describe the modelled species at steady-state:

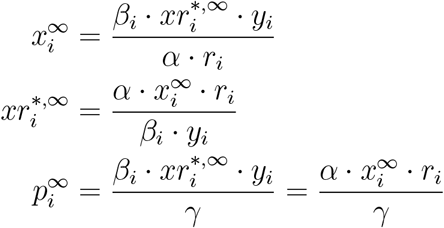

The effective initiation and elongation rates measure the net contribution of the initiation and elongation steps, respectively, and are defined as:

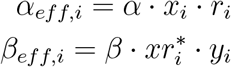

According to Equations 1 and 2 and the conditions of the steady-state

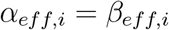

at steady-state.

### Measuring competition

For measuring the competition between long and short mRNA protein synthesis at equilibrium, we define *ρ* as the change of the mean protein yield ratios.

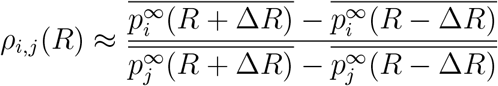

This metric is an approximation of the gradient of the long-to-short protein ratios with regards to ribosome abundance. The long-to-short protein ratio is defined as:

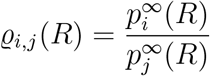

While this ratio informs whether there occurs a length-dependent shift in the protein yield, it is not able to identify the direction of this shift. The protein yield is strictly positive *p*_*i*_ *>* 0*∀i*. It follows, *ϱ*_*i,j*_(*R*) *>* 0. A reduced ratio can be caused by an increase in the denominator or a reduction of the numerator. To disambiguate between these two cases, we calculated the ratio of the approximated gradient of the protein yield *ρ*_*i,j*_ with regard to ribosome abundance *R*:

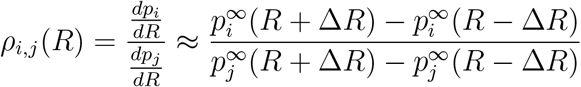

Using this quantity, we can distinguish two conditions:

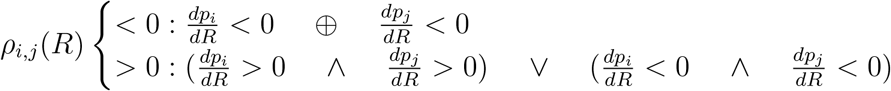

Thus, the competition measurement *ρ*_*i,j*_ *<* 0 only if the signs of the gradient of long and short proteins are different. If both long and short proteins are upregulated or downregulated, *ρ*_*i,j*_ *>* 0. If *i* and *j* denote groups of proteins, e.g. all proteins longer than 2*k i* : {*x*| *x >* 2*k*} and all proteins shorter than 2*k j* :{ *x*| *x <* 2*k*}, the gradient is approximated over the mean protein yields of the respective subpopulations:

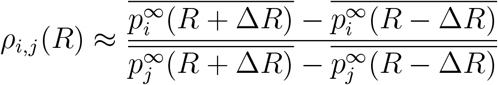

### Reproducibility audit

A reproducibility audit^31^ tests the feasibility of project code for reruns. Conducting such an audit ensures that any provided code can be run on a private machine with minimal knowledge of the model or project details. It ensures that the results can be reproduced by others independent of the specific set-up.

The available code underwent a reproducibility audit in two rounds. A round was counted as complete after a discussion of any undisclosed dependencies and encountered run time errors. We assessed whether the code was executable without any prior knowledge of the model specification nor its neurological background. A detailed description of the round-specific feedback from the reproducibility audit is provided in the Appendix.

## Code availability

All code is available at https://github.com/prubisch/competition_in_proteinsynthesis_in_autism.

## ACKNOWLEDGMENTS

J.A.H.W. was funded by the BMFTR, grant number 01ZZ2019 (MeDaX). P.R. was funded by an EPSRC DTA scholarship. S.B.R. was funded by a grant from the Biotechnology and Biological Sciences Research Council for the South West Biosciences Doctoral Training Partnership (DTP3: BB/T008741/1)

We thank Isabel Cornacchia for her helpful comments on earlier presentations of this project.

## AUTHOR CONTRIBUTIONS

P.R.: conceptualization, methodology, investigation, formal analysis, visualisation, software, writing - original draft, review & editing

S.B.R.: conceptualization, writing - review & editing

J.A.H.W.: funding acquisition, validation, software, writing review & editing

M.I.S.: conceptualization, methodology, formal analysis, resources, supervision, writing - review & editing

C.D.: conceptualization, formal analysis, supervision, writing - review & editing

## Appendix

### Generalisation to randomly drawn populations

The results shown in the main paper are derived considering one randomly drawn population. To demonstrate that the results generalise to multiple realisations, we drew 100 random populations and computed the mean (solid line), median (dotted line) and standard deviation of the protein yield. The results are shown in Figure 7 and are equivalent to the results shown in 3A of the main text. They demonstrate that the exact elongation rates become increasingly significant at hihger ribosomal abundances. Especially, the reversion point between initiation- and elongation- restricted regime denotes an increase in importance of exact elongation rates shown by the increase in the standard deviation (7B: Competitive R).

**Figure 7.**
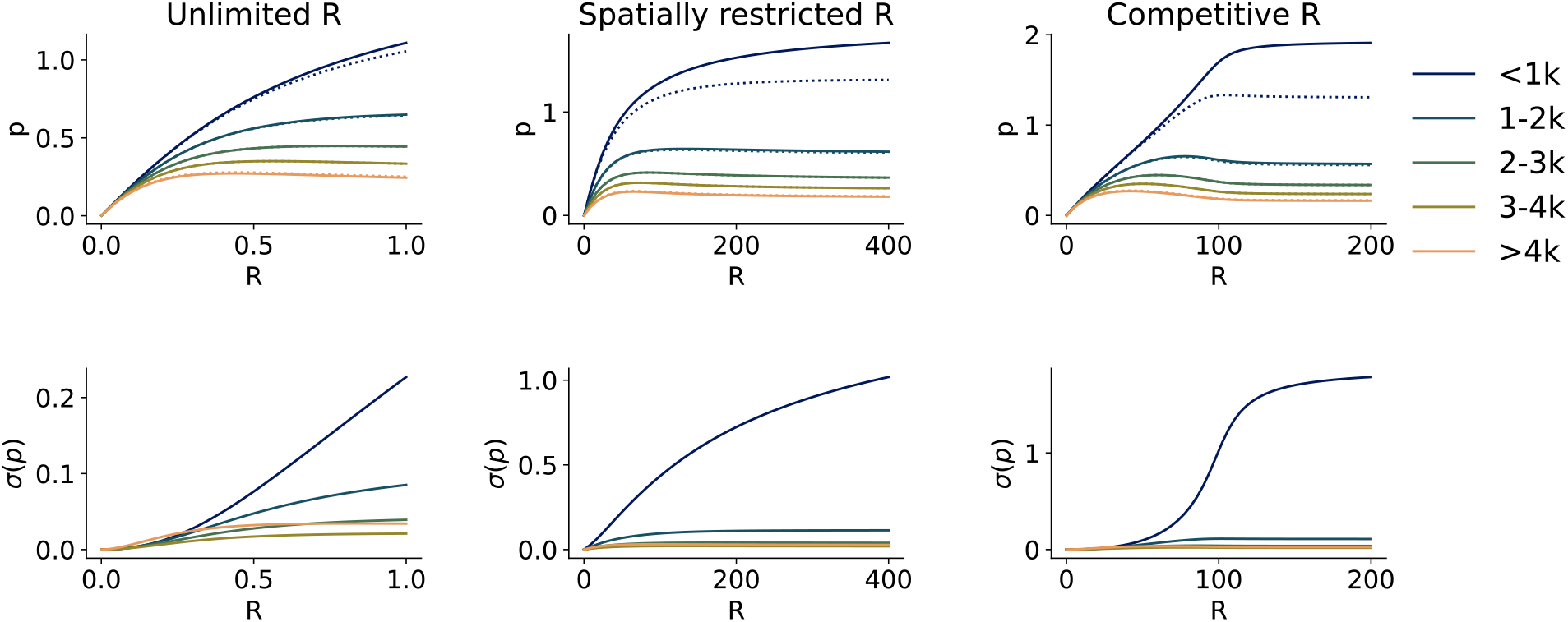
Statistics of the protein yield of 100 population realisations according to the amino-acid competitive models. **Top:** Mean (solid) and median (dotted) protein yield of the subpopulations according to length. **Bottom:** Standard deviation of the protein yield at equilibrium.

### Reproducibility audit

The following sections lists all problems identified in the reproducibility audit and how they were addressed. Between the first and the second round the auditor met with the main code developer in person to discuss the results in detail and propose bigger changes. The auditor were not given any specific prompts before carrying out the audit other than “Please reproduce the figures in the preprint with the provided information in the repository.”.

#### Round 1

~~~
 
 •README.md
     
    ‐ typos : corrected spelling and added additional information
 
 •figure1_mock_data.ipynb
   ‐ not able to import figures_helper.py : restarting kernel (no code changes)
   ‐ FileNotFound Error: figures/fig1_problem_description.pdf : directory check and mkdir for figures/
 • figure2_baysian_optimisation_all_models.ipynb
 
  ‐ ValueError in run_opt(): notebook no longer in use, analysis in Figure 2 was fundamentally reworked between audits
 • figure3_to_6_simulation_all_model_variations.ipynb
  
 ‐ Error: No module named ‘cmcrameri’ : added module cmcrameri including module version to prerequisite list in README.md
  ‐ missing directory simulation_results/: directory check and mkdir for
simulation_results/
  ‐ figures not saved in /figures/: save_fig=True in plotting function calls
  ‐ typos in comments: corrected spelling
~~~

#### In-person discussion between round 1 and round 2

The accessibility of jupyter notebooks was discussed. Not all universities provide an environment in which cloning and running jupyter notebooks including non-standard module imports is possible without either dedicated server time or the installation of needed packages through the IT service. Therefore, all jupyter notebooks were moved into a dedicated folder and replaced with python scripts in the main directory. The decision to still supply the jupyter notebooks has been made to provide interested researchers with a possible start point for further development outside of the projects repository. To reproduce the results, the python scripts are sufficient. They prompt the user to select options during the run process, e.g. selecting whether rerunning the simulations with new parameters or recreating the figures from the provided data. System checks for subdirectories have been included. Should these checks fail due to an incomplete set-up or only partial cloning of the main-branch, subdirectories are established explicitly.

#### Round 2

~~~
• README.md
   ‐ sklearn Deprecation Warning:corrected to scikit-learn package (equivalent without deprecation warning)
• figure1_mock_data.py and figure1_mock_data.ipynb
  ‐ OSError: ‘seaborn-colorblind’ is not a valid package style in plotting_helper.py: Solution was already provided, second run was successful, lines 9-12 in plotting_helper.py pertaining to matplotlib.style and seaborn-colorblind were deleted as colour styles are specified using cmcrameri package making them superfluous
• figure2_parameter_exploration_models.py and
figure2_parameter_exploration_models.ipynb
  ‐ ran successful
• figure3_to_6_simulation_all_model_variations.py and
figure3_to_6_simulation_all_model_variations.ipynb
 
  ‐ ran successful
~~~

## References

1. Sun, C., and Schuman, E. M. (2022). Logistics of neuronal protein turnover: Numbers and mechanisms. Molecular and Cellular Neuroscience 123. doi:10.1016/j.mcn.2022.103793.

2. Holt, C. E., Martin, K. C., and Schuman, E. M. (2019). Local translation in neurons: visualization and function. Nature Structural & Molecular Biology 26. doi:10.1038/s41594-019-0263-5.

3. Jiang, C., and Schuman, E. M. (2002). Regulation and function of local protein synthesis in neuronal dendrites. Trends in Biochemical Sciences 27. doi:10.1016/S0968-0004(02)02190-4.

4. Sutton, M. A., and Schuman, E. M. (2006). Dendritic Protein Synthesis, Synaptic Plasticity, and Memory. Cell 127. doi:10.1016/j.cell.2006.09.014.

5. Fonseca, R., Vabulas, R. M., Hartl, F. U., Bonhoeffer, T., and Nägerl, U. V. (2006). A Balance of Protein Synthesis and Proteasome-Dependent Degradation Determines the Maintenance of LTP. Neuron 52. doi:10.1016/j.neuron.2006.08.015.

6. Ostroff, L. E., Fiala, J. C., Allwardt, B., and Harris, K. M. (2002). Polyribosomes Redistribute from Dendritic Shafts into Spines with Enlarged Synapses during LTP in Developing Rat Hippocampal Slices. Neuron 35. doi:10.1016/S0896-6273(02)00785-7.

7. Sun, C., Nold, A., Fusco, C. M., Rangaraju, V., Tchumatchenko, T., Heilemann, M., and Schuman, E. M. (2021). The prevalence and specificity of local protein synthesis during neuronal synaptic plasticity. Science Advances 7. doi:10.1126/sciadv.abj0790.

8. Chen, P. B., Kawaguchi, R., Blum, C., Achiro, J. M., Coppola, G., O’Dell, T. J., and Martin, K. C. (2017). Mapping Gene Expression in Excitatory Neurons during Hippocampal Late-Phase Long-Term Potentiation. Frontiers in Molecular Neuroscience 10. doi:10.3389/fnmol.2017.00039.

9. Hou, L., Antion, M. D., Hu, D., Spencer, C. M., Paylor, R., and Klann, E. (2006). Dynamic Translational and Proteasomal Regulation of Fragile X Mental Retardation Protein Controls mGluR-Dependent Long-Term Depression. Neuron 51. doi:10.1016/j.neuron.2006.07.005.

10. Hien, A., Molinaro, G., Liu, B., Huber, K. M., and Richter, J. D. (2020). Ribosome profiling in mouse hippocampus: plasticity-induced regulation and bidirectional control by TSC2 and FMRP. Molecular Autism 11. doi:10.1186/s13229-020-00384-9.

11. Besse, F., and Ephrussi, A. (2008). Translational control of localized mRNAs: restricting protein synthesis in space and time. Nature Reviews Molecular Cell Biology 9. doi:10.1038/nrm2548.

12. Kang, H., and Schuman, E. M. (1996). A Requirement for Local Protein Synthesis in Neurotrophin-Induced Hippocampal Synaptic Plasticity. Science 273. doi:10.1126/science.273.5280.1402.

13. Schimanski, L. A., and Barnes, C. A. (2010). Neural Protein Synthesis during Aging: Effects on Plasticity and Memory. Frontiers in Aging Neuroscience 2. doi:10.3389/fnagi.2010.00026.

14. Tao, J., Wu, H., Coronado, A. A., de Laittre, E., Osterweil, E. K., Zhang, Y., and Bear, M. F. (2016). Negative Allosteric Modulation of mGluR5 Partially Corrects Pathophysiology in a Mouse Model of Rett Syndrome. The Journal of Neuroscience 36. doi:10.1523/JNEUROSCI.0672-16.2016.

15. Fusco, C. M., Desch, K., Dörrbaum, A. R., Wang, M., Staab, A., Chan, I. C. W., Vail, E., Villeri, V., Langer, J. D., and Schuman, E. M. (2021). Neuronal ribosomes exhibit dynamic and context-dependent exchange of ribosomal proteins. Nature Communications 12. doi:10.1038/s41467-021-26365-x.

16. Genuth, N. R., and Barna, M. (2018). The Discovery of Ribosome Heterogeneity and Its Implications for Gene Regulation and Organismal Life. Molecular Cell 71. doi:10.1016/j.molcel.2018.07.018.

17. Mills, E. W., and Green, R. (2017). Ribosomopathies: There’s strength in numbers. Science 358. doi:10.1126/science.aan2755.

18. Farley-Barnes, K. I., Ogawa, L. M., and Baserga, S. J. (2019). Ribosomopathies: Old Concepts, New Controversies. Trends in Genetics 35. doi:10.1016/j.tig.2019.07.004.

19. Seo, S. S., Louros, S. R., Anstey, N., Gonzalez-Lozano, M. A., Harper, C. B., Verity, N. C., Dando, O., Thomson, S. R., Darnell, J. C., Kind, P. C., Li, K. W., and Osterweil, E. K. (2022). Excess ribosomal protein production unbalances translation in a model of fragile x syndrome. Nature Communications 13. doi:10.1038/s41467-022-30979-0.

20. Witte, F., Ruiz-Orera, J., Mattioli, C. C., Blachut, S., Adami, E., Schulz, J. F., SchneiderLunitz, V., Hummel, O., Patone, G., Mücke, M. B., Šilhavý, J., Heinig, M., Bottolo, L., San-chis, D., Vingron, M., Chekulaeva, M., Pravenec, M., Hubner, N., and Van Heesch, S. (2021). A trans locus causes a ribosomopathy in hypertrophic hearts that affects mRNA translation in a protein length-dependent fashion. Genome Biology 22. doi:10.1186/s13059-021-02397-w.

21. Louros, S. R., Seo, S. S., Maio, B., Martinez-Gonzalez, C., Gonzalez-Lozano, M. A., Muscas, M., Verity, N. C., Wills, J. C., Li, K. W., Nolan, M. F., and Osterweil, E. K. (2023). Excessive proteostasis contributes to pathology in fragile X syndrome. Neuron 111. doi:10.1016/j.neuron.2022.11.012.

22. Darnell, J. C., VanbDriesche, S. J., Zhang, C., Hung, K. Y. S., Mele, A., Fraser, C. E., Stone, E. F., Chen, C., Fak, J. J., Chi, S. W., Licatalosi, D. D., Richter, J. D., and Darnell, R. B. (2011). FMRP Stalls Ribosomal Translocation on mRNAs Linked to Synaptic Function and Autism. Cell 146. doi:10.1016/j.cell.2011.06.013.

23. McGirr, T., Onar, O., and Jafarnejad, S. M. (2024). Dysregulated ribosome quality control in human diseases. The FEBS Journal 292. doi:10.1111/febs.17217.

24. Scarpitti, M. R., Pastore, B., Tang, W., and Kearse, M. G. (2024). Characterization of ribosome stalling and no-go mRNA decay stimulated by the fragile X protein, FMRP. Journal of Biological Chemistry 300. doi:10.1016/j.jbc.2024.107540.

25. Kute, P. M., Labun, K., Tjeldnes, H., Valen, E., and Muddashetty, R. (2026). Ribosome profiling to study translation in cortical synaptoneurosomes on group 1 mGluR activation. bioRxiv. doi:10.64898/2026.05.10.723250.

26. Lodish, H. F. (1974). Model for the regulation of mRNA translation applied to haemoglobin synthesis. Nature 251. doi:10.1038/251385a0.

27. Raveh, A., Margaliot, M., Sontag, E. D., and Tuller, T. (2016). A model for competition for ribosomes in the cell. Journal of The Royal Society Interface 13. doi:10.1098/rsif.2015.1062.

28. Dao Duc, K., and Song, Y. S. (2018). The impact of ribosomal interference, codon usage, and exit tunnel interactions on translation elongation rate variation. PLOS Genetics 14. doi:10.1371/journal.pgen.1007166.

29. Rogers, D. W., Böttcher, M. A., Traulsen, A., and Greig, D. (2017). Ribosome reinitiation can explain length-dependent translation of messenger RNA. PLOS Computational Biology 13. doi:10.1371/journal.pcbi.1005592.

30. Fernandes, L. D., de Moura, A. P., and Ciandrini, L. (2017). Gene length as a regulator for ribosome recruitment and protein synthesis: theoretical insights. Scientific Reports 7. doi:10.1038/s41598-017-17618-1.

31. Fitzpatrick, R., and Stefan, M. I. (2022). Validation Through Collaboration: Encouraging Team Efforts to Ensure Internal and External Validity of Computational Models of Biochemical Pathways. Neuroinformatics 20. doi:10.1007/s12021-022-09584-5.

